# Do nanoplastics reshape microglial support of neuronal resilience? A study of microglial bioenergetics and microglia–neuron communication *in vitro*

**DOI:** 10.64898/2026.06.17.732827

**Authors:** Electra Brunialti, Alessandro Villa, Clara Meda, Parolini Marco, Paolo Ciana, Lavina Casati

## Abstract

Nanoplastics (NPs) are emerging environmental contaminants able to cross biological barriers, disrupt cellular and organelle homeostasis, and alter the brain microenvironment. This study investigated whether NPs affect microglia–neuron communication, a key mechanism underlying neuronal resilience, via the nuclear factor erythroid 2-like 2 (NFE2L2) pathway. Using an *in vitro* model, we evaluated the effects of polystyrene nanoplastics on microglial metabolic fitness and microglia-mediated neuronal stress responses. Increasing NP concentrations induced a dose-dependent biphasic effect. Low to intermediate concentrations increased intracellular adenosine triphosphate (ATP) levels in microglia and enhanced microglia-mediated activation of neuronal NFE2L2. In contrast, high NP concentration impaired microglial metabolism, reduced ATP availability, and decreased microglia–neuron communication. These findings indicate that NPs alter microglial energetic status and modulate neuroprotective signalling, potentially contributing to impaired neuron–microglia interactions and increased susceptibility to neurotoxicity.

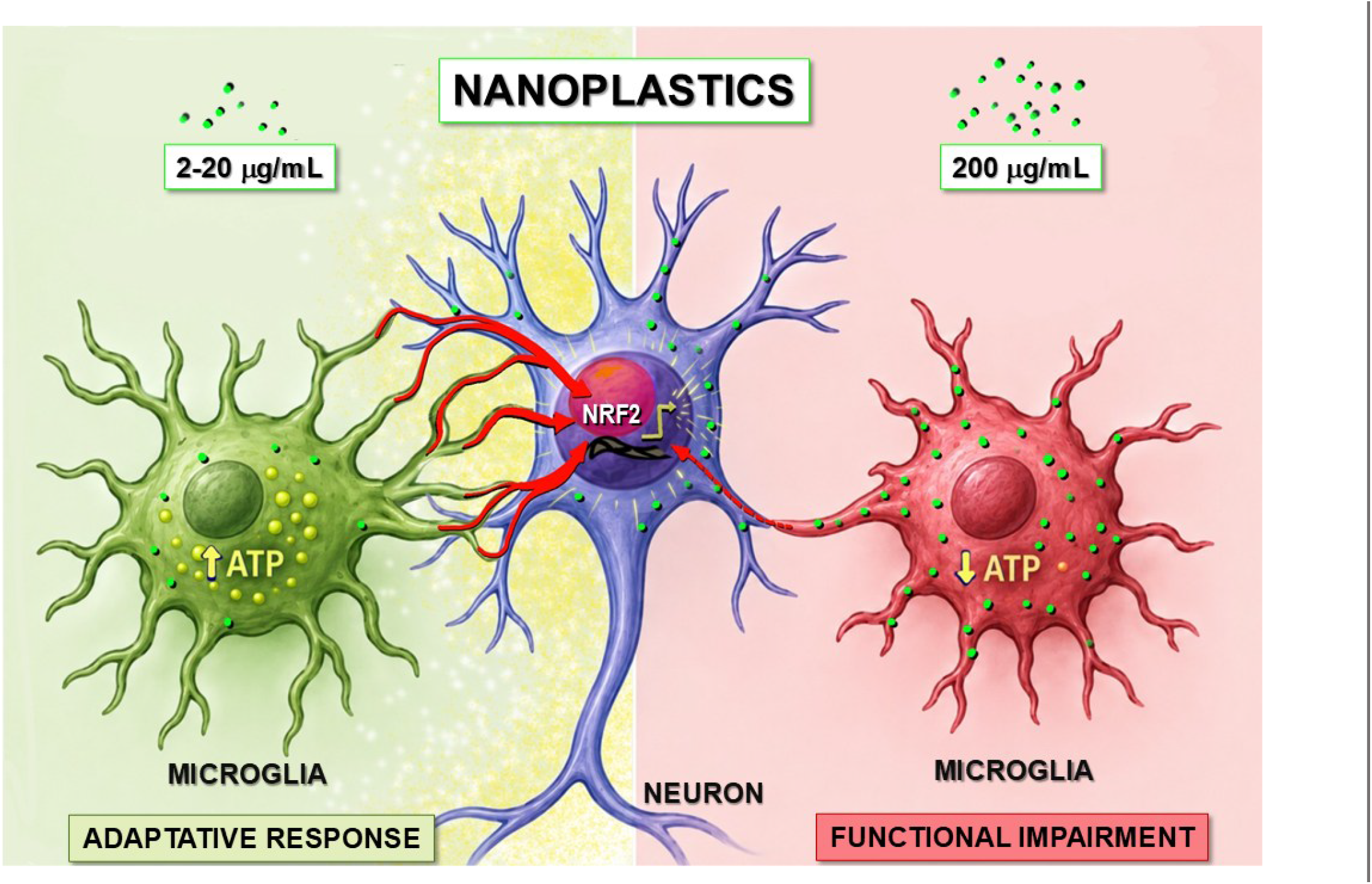

**Highlights:**

1. Nanoplastics alter microglial metabolic fitness *in vitro*.
2. Nanoplastics biphasically modulate microglial support of neurons.
3. High nanoplastic concentration reduces microglial support to neurons.
4. Microglial bioenergetics may link nanoplastics to neuronal vulnerability.

## 1. Introduction

Nanoplastics (NPs) are emerging environmental contaminants of increasing concern for human health. Growing evidence indicates that NPs can accumulate in human tissues—including the placenta, brain, bone marrow, macrophages, and atherogenic plaques—and are capable of crossing biological barriers and membranes (Nihart et al., 2025, de Sousa et al., 2024, Bianchi et al., 2025, Marfella et al., 2024, Guo et al., 2024, Jani et al., 1990, Jing et al., 2022). At the cellular level, NPs are readily internalized and can disrupt fundamental cellular functions, including intercellular communication (Villa et al., 2026, Sangkham et al., 2022, Yee et al., 2021).

NPs interfere with mitochondrial function, a central hub for cellular metabolism and signalling. Nanomaterial-induced mitochondrial dysfunction has been associated with impaired energy production, altered bioenergetic states, and disruption of intracellular trafficking pathways (Lima et al., 2022, Wu et al., 2020). In this context, NPs have recently emerged as disruptors of mitochondrial integrity and intracellular communication (Jain and Zoncu, 2022). At the mitochondrial level, NPs impair membrane potential and alter mitochondrial dynamics, resulting in defective oxidative phosphorylation and reduced ATP production, thereby compromising cellular energy homeostasis (Maharana et al., 2026). This energetic imbalance is accompanied by increased generation of reactive oxygen species (ROS) (Giannandrea et al., 2024, Li et al., 2023, De Felice et al., 2022), leading to oxidative stress that damages mitochondrial DNA, lipids, and proteins (Maharana et al., 2026).

In parallel, NPs dysregulate calcium signalling and redox homeostasis, further exacerbating mitochondrial dysfunction and activating stress-response pathways, including mitophagy and apoptosis. Importantly, mitochondrial impairment extends beyond bioenergetics, affecting mitochondria’s ability to coordinate signalling with other organelles, such as the endoplasmic reticulum and the nucleus, thereby disrupting intracellular communication networks (Trevisan et al., 2019, Xu et al., 2023, Maharana et al., 2026). Alterations in mitochondrial dynamics, including fusion, fission, and motility, have been associated with neurological disorders, cancer, and metabolic diseases (Chen et al., 2023a, Chen et al., 2023b).

These processes are particularly relevant in the central nervous system, where mitochondrial function plays a critical role in neuron–microglia communication (Brunialti et al., 2021, Brunialti et al., 2025). Microglia rely on tightly regulated metabolic programs and mitochondrial activity to sustain essential functions such as surveillance, motility, and neuroprotection. Perturbations in mitochondrial function and cellular energy balance can therefore significantly impact microglial physiology and their ability to support neuronal homeostasis.

In our previous work, we demonstrated that microglia, through direct physical interactions with neurons, promote neuronal detoxification by modulating the NFE2L2 (also known as NRF2) pathway, a key regulator of oxidative stress responses (Brunialti et al., 2021). Notably, this regulatory mechanism is closely linked to cellular metabolism and mitochondrial function. Furthermore, impairment of microglial function induced by environmental or metabolic stressors increases neuronal susceptibility to pathological phenotypes (Brunialti et al., 2025).

Building on this framework and considering emerging evidence that mitochondrial dysfunction and ATP depletion critically affect microglial function, this study investigated whether NPs impair microglial neuroprotective functions. Specifically, we assessed whether NPs alter microglial energetic metabolism and compromise neuron– microglia communication, thereby potentially contributing to loss of cellular homeostasis.

## 2. Methods

### Cell culture and co-culture conditions

All cell lines were obtained from the American Type Culture Collection (ATCC). SK-ARE-*luc2* cells were generated by stable transfection of SK-N-BE neuroblastoma cells with the pARE-luc2-ires-tdTomato reporter plasmid (Rizzi et al., 2018). For co-culture experiments, SK-ARE-*luc2* cells were seeded at 60,000 cells/well in 24-well plates and allowed to adhere for 24 h. BV-2 microglial cells were then seeded on top of the neuronal layer at a density of 7,000 cells/well. For BV-2 monoculture experiments, cells were seeded at a density of 25,000 cells/well. Cells were maintained in Neurobasal-A medium (Cat. 10888-022, Life Technologies) supplemented with 1% penicillin–streptomycin, 1% GlutaMAX (Cat. 35050061, Life Technologies), 2% B-27 Supplement (Cat. 17504-044, Gibco), and 10 mM HEPES (Cat. H0887, Merck). Cultures were kept at 37 °C in a humidified atmosphere containing 5% CO_2_.

### Nanoplastic exposure and ATP stimulation

Cells were exposed for 48 h to green fluorescent nanoplastics (NPs; nominal diameter 0.047 μm; Cat. G50, Life Technologies) at increasing concentrations of 0, 2, 20, and 200 μg/mL. Vehicle-treated cells received the corresponding volume of PBS. Where indicated, ATP (Cat. A7699, Merck) was administered at a final concentration of 50 μM for 4 h after 48 h of NP exposure. Water was used as a vehicle control for ATP treatment.

### Imaging acquisition

Bright-field and fluorescence images were acquired using an Axiovert 200M microscope equipped with AxioVision software (Rel. 4.9, Zeiss). For each experimental condition, 10 random fields were acquired using bright-field imaging and the green fluorescence channel to detect fluorescent nanoplastics. Images were subsequently processed using Fiji software (ImageJ, NIH, version 2.0.0).

### Luciferase activity assay

Luciferase activity was measured as previously described (Brunialti et al., 2021). Briefly, cells were lysed using Luciferase Cell Culture Lysis Reagent (Cat. E1531, Promega). Luciferase activity was then measured using luciferase assay buffer, and luminescence emission was recorded with a Veritas luminometer (Turner Biosystems). Relative luminescence units (RLU) were determined and used as a readout of NFE2L2/ARE reporter activation.

### Intracellular ATP quantification

Intracellular ATP levels were measured using the Luminescent ATP Detection Assay Kit (Cat. ab113849, Abcam), according to the manufacturer’s instructions. Bioluminescence was measured using a Veritas luminometer (Turner Biosystems), and ATP concentrations were determined using a standard calibration curve.

### MTT assay

Cellular reductive metabolic activity was assessed using the MTT assay. Briefly, MTT reagent (Cat. 475989, Merck) was added directly to the culture medium at a final concentration of 0.5 mg/mL. After 30 min of incubation at 37 °C, cells were lysed with DMSO (Cat. D8418, Merck), and the absorbance at 595 nm was measured. Data were expressed as the percentage of substrate conversion relative to vehicle-treated cells.

### Protein quantification

Total protein content was measured using the Bradford assay (Cat. 23238, Thermo Scientific) according to the manufacturer’s instructions.

### Statistical analysis

Data are presented as mean ± standard error of the mean (SEM). Statistical analyses were performed using GraphPad Prism version 8.0 (GraphPad Software Inc.). Statistical outliers were identified and excluded using the ROUT method with Q = 1. One-way ANOVA was used for comparisons among three or more independent groups. Two-way ANOVA was used to assess the effects of two independent variables. Post-hoc tests are indicated in the corresponding figure legends. A p-value < 0.05 was considered statistically significant.

## 3. Results

### Nanoplastic exposure modulates microglial metabolism and microglia–neuron communication

To investigate the effects of NPs on microglia–neuron interactions, we used SK-N-BE neuroblastoma cells as a dopaminergic-like neuronal model together with immortalized BV-2 microglial cells.

Fluorescent polystyrene NPs with a nominal diameter of 0.05 μm were used as a representative NPs model (Giannandrea et al., 2024, Villa et al., 2026). Cells were exposed to increasing NP concentrations, 0, 2, 20, and 200 μg/mL, for 48 h; a range selected to cover a broad *in vitro* exposure window, from low/intermediate doses to a high-concentration condition. The 48-h exposure time was chosen as a commonly used window suitable for assessing NP–cell interaction, potential cellular accumulation, and the development of detectable cellular responses (Giannandrea et al., 2024).

Preliminary imaging analysis showed that NP-associated fluorescence was mainly detected within the cellular area, suggesting close association with, or uptake by, exposed cells (Figure 1A). No evident morphological alterations were observed in either BV-2 or SK-N-BE cells at any concentration tested (Figure 1B). Consistently, total protein content remained unchanged across experimental conditions (Figure 1C), suggesting the absence of overt cytotoxicity.

**Figure 1.**
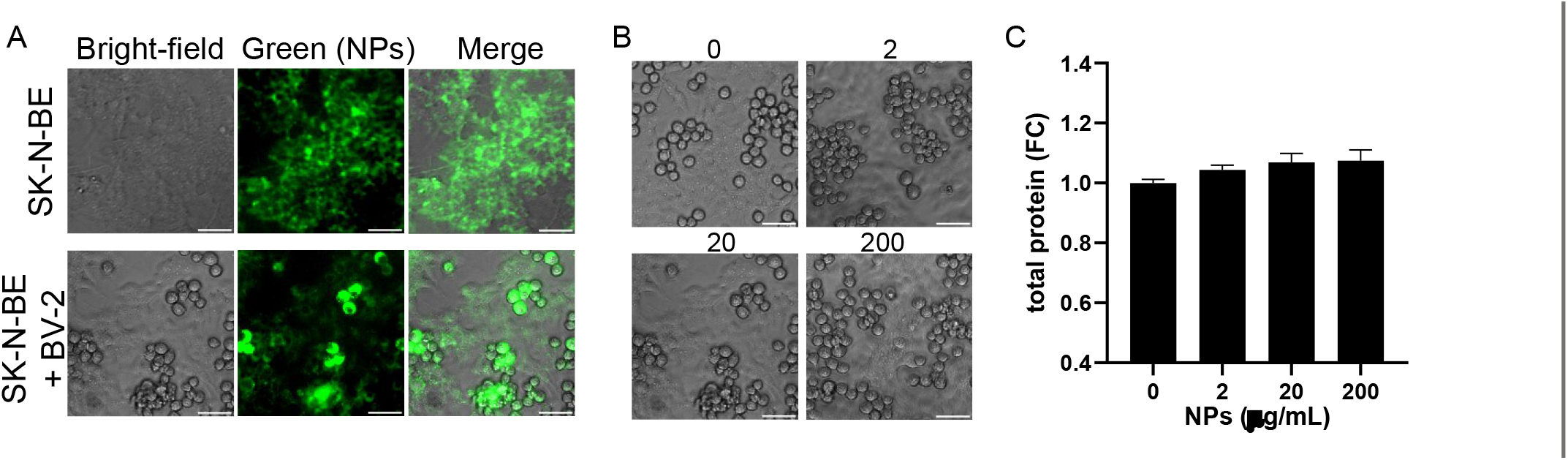
General assessment of nanoplastics-treated cultures. **(A)** Representative images of SK-N-BE cells exposed to 20 μg/mL nanoplastics (NPs) for 48 h. Bright-field images (left), NP fluorescence signal (green channel, middle), and merged images (right) are shown. Scale bar = 50 μm. **(B)** Representative bright-field images of SK-N-BE/BV-2 co-cultures following 48 h exposure to increasing NP concentrations (0–200 μg/mL). **(C)** Total protein content in co-cultures following 48 h NPs exposure (0–200 μg/mL). Data are expressed as mean ± SEM (n = 12). No statistically significant differences were detected among groups (one-way ANOVA).

To evaluate whether NPs affect the functional interaction between microglia and neuronal cells, we employed a previously established co-culture system in which SK-N-BE cells stably carrying the ARE- *luc2* reporter construct, hereafter referred to as SK-ARE-*luc2* cells, were cultured together with BV-2 microglia (Figure 2A) (Brunialti et al., 2021). In this model, the neuronal bioluminescent signal reflects NFE2L2-dependent transcriptional activity and provides a functional readout of BV-2-mediated modulation of the neuronal NFE2L2 pathway, used as an index of microglia-to-neuron communication.

**Figure 2.**
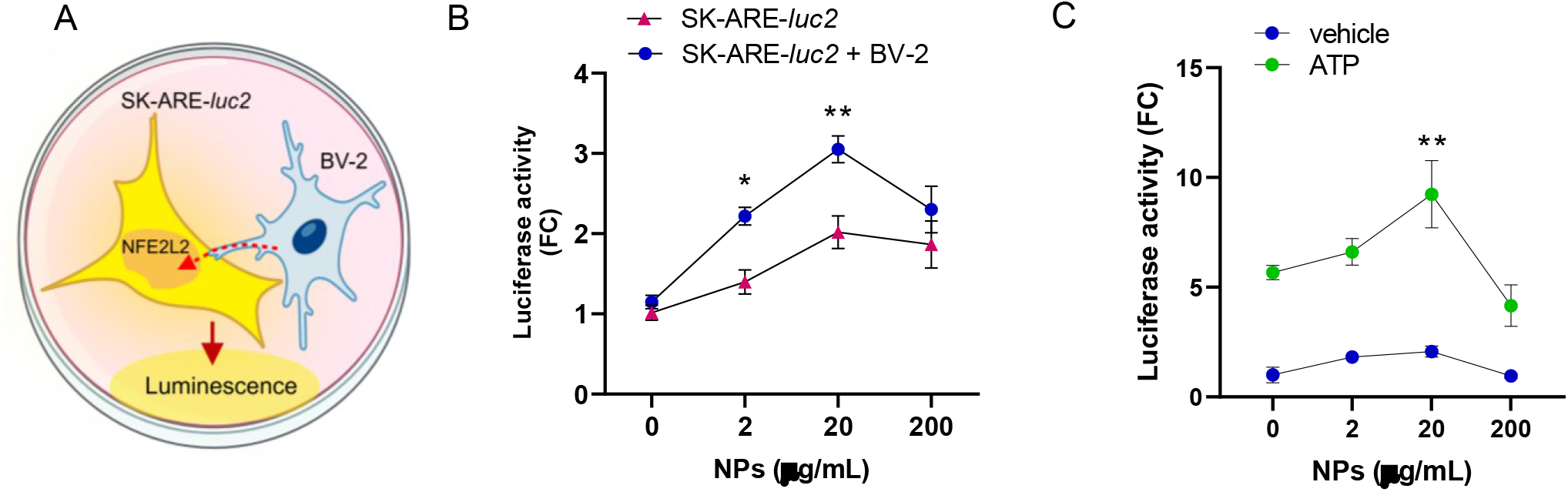
Nanoplastics modulate microglia–neuron communication. **(A)** Schematic representation of the co-culture system used to assess microglia–neuron communication. BV-2 microglia promote neuronal activation of the NFE2L2 (NRF2) pathway, which is detected using SK-ARE-*luc2* reporter cells. The resulting bioluminescent signal provides a semi-quantitative measure of microglia-to-neuron communication. **(B)** SK-ARE-*luc2* cells alone or co-cultured with BV-2 cells were exposed to increasing concentrations of nanoplastics (NPs; 0–200 μg/mL) for 48 h. Luciferase activity is expressed as fold change (FC) versus vehicle-treated SK-ARE-*luc2* cells and presented as mean ± SEM (n = 6). *p < 0.05, **p < 0.01 versus SK-ARE-*luc2* cells at the same NP concentration (two-way ANOVA, Sidak’s test). **(C)** SK-ARE-*luc2*/BV-2 co-cultures were exposed to NPs for 48 h and stimulated with ATP (50 μM, 4 h). Luciferase activity are expressed as FC versus vehicle-treated cultures and presented as mean ± SEM (n = 6). **p < 0.01 indicates differences versus the corresponding 0 μg/mL NP condition within each treatment group, vehicle or ATP-treated cells (two-way ANOVA, Dunnett’s test).

Exposure to NPs increased NFE2L2-related luciferase activity in SK-ARE-*luc2* cells, with the response rising up to 20 μg/mL and reaching a plateau between 20 and 200 μg/mL (Figure 2B). These data indicate activation of the neuronal NFE2L2 pathway and are consistent with previous observations reported in other experimental settings (Gettings et al., 2026). In co-culture conditions, the BV-2-dependent modulation of neuronal NFE2L2 activity displayed a biphasic response following NP exposure. At low and intermediate concentrations (2 and 20 μg/mL), NPs significantly enhanced the co-culture-associated communication signal. Conversely, exposure to 200 μg/mL markedly reduced this signal, suggesting a substantial loss of the BV-2-mediated contribution to neuronal NFE2L2 activation. These findings suggest that low NP concentrations may trigger adaptive or compensatory responses, whereas higher concentrations may impair microglia’s ability to sustain functional support for neuronal cells.

To further investigate the responsiveness of the co-culture system, cells were stimulated with 50 μM ATP for 4 h, following NP exposure (Figure 2C) or with vehicle (water) as a control. ATP is known to activate purinergic signalling and has been associated with increased oxidative phosphorylation in microglial cells and with promoting microglia–neuron communication (Brunialti et al., 2025, Hu et al., 2020, Ledderose et al., 2020). As expected, ATP induced an approximately five-fold increase in communication signal in cultures not exposed to NPs. This response was further enhanced in co-cultures treated with low (2 μg/mL) and intermediate (20 μg/mL) NP concentrations. Conversely, following exposure to 200 μg/mL NPs, ATP-induced communication was markedly blunted, decreasing to approximately a 4-fold increase compared with the 9-fold response observed at 20 μg/mL. These findings suggest that high NP concentrations reduce the co-culture system’s ability to respond to ATP stimulation, possibly reflecting altered purinergic-dependent microglial signalling and reduced ATP-mediated functional interactions between microglia and neuronal cells.

Since we previously showed that efficient microglia–neuron communication requires preserved microglial metabolic fitness (Brunialti et al., 2025), the changes in the communication signal observed after NPs exposure prompted us to investigate whether NPs induce metabolic and energetic alterations in microglial cells. To this purpose, we quantified intracellular ATP levels in BV-2 cells, as ATP is a key energy-producing molecule whose reduction has been associated with impaired microglia–neuron communication (Brunialti et al., 2025). Microglial ATP content increased by approximately 10% at 2 and 20 μg/mL NPs, paralleling the enhancement of the co-culture communication signal observed under these conditions (Figure 2B). In contrast, ATP levels were reduced by approximately 10% following exposure to 200 μg/mL NPs, closely mirroring the loss of communication efficiency observed at the same concentration (Figure 2B). Together, these observations suggest that NP-induced alterations in microglia–neuron communication may be at least partly linked to energetic remodelling in microglial cells.

Since ATP levels reflect cellular energetic availability but do not fully capture the overall metabolic state of microglial cells, we next assessed MTT reduction to evaluate whether NP exposure also affected cellular reductive metabolic activity (Figure 3B). MTT conversion in BV-2 cells was reduced across the entire NP concentration range, with a comparable decrease of approximately 10% between 2 and 200 μg/mL, suggesting an early and persistent alteration of microglial reductive metabolism.

**Figure 3.**
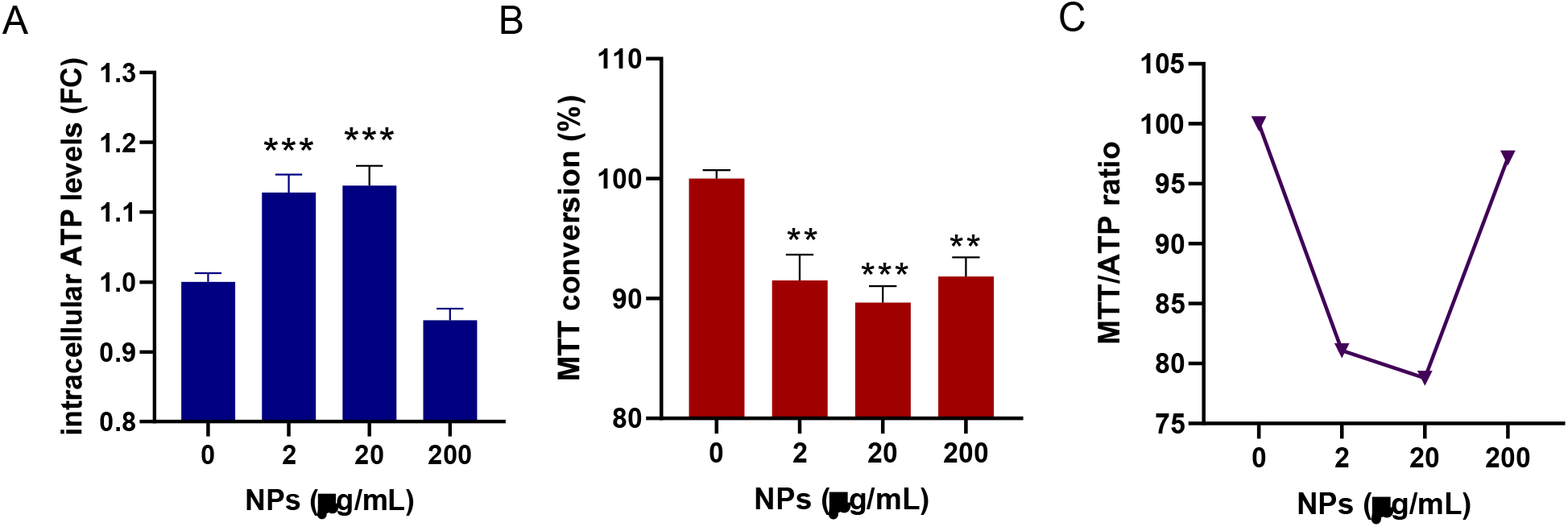
Nanoplastics affect microglial metabolic fitness. BV-2 cells were exposed to increasing concentrations of nanoplastics (NPs; 0–200 μg/mL) for 48 h. **(A)** Intracellular ATP levels. Data are expressed as fold change (FC) relative to vehicle-treated cells and presented as mean ± SEM (n = 12). **p < 0.01, ***p < 0.001 versus vehicle-treated cells (one-way ANOVA followed by Dunnett’s multiple comparisons test). **(B)** MTT reduction assay. Data are expressed as percentage of substrate conversion relative to vehicle-treated cells and presented as mean ± SEM (n = 6). **p < 0.01, ***p < 0.001 versus vehicle-treated cells (one-way ANOVA followed by Dunnett’s multiple comparisons test). **(C)** MTT/ATP ratio calculated from the MTT conversion values shown in panel B and the intracellular ATP levels shown in panel A.

To further explore the relationship between cellular reductive metabolism and ATP availability, we calculated the MTT/ATP ratio as an exploratory descriptive index (Figure 3C). MTT reduction reflects cellular reducing capacity, whereas intracellular ATP levels provide an estimate of energy availability.

The MTT/ATP ratio decreased at 2 and 20 μg/mL, indicating that the increase in ATP content was not accompanied by a proportional increase in MTT conversion. This divergent behaviour suggests a selective remodelling of microglial metabolism at low and intermediate NP concentrations. Conversely, at 200 μg/mL, the ratio increased as ATP levels declined, suggesting a shift from an adaptive metabolic response toward energetic impairment. Overall, the opposite behaviour at low/intermediate doses and at the highest dose supports the presence of dose-dependent metabolic remodelling rather than a uniform decrease in cell viability.

## 4. Discussion

Micro- and nanoplastics are pervasive environmental contaminants detected in air, soil, water, food, and human biological samples, raising increasing concern about their potential impact on human health(Ali et al., 2025). Because of their small size, NPs may interact with biological barriers, enter cells, and accumulate within intracellular compartments, including endo-lysosomal structures.(Maharana et al., 2026) Experimental studies have linked NP exposure to oxidative stress, mitochondrial dysfunction, lysosomal impairment, and activation of innate immune pathways, all of which are highly relevant to brain homeostasis(Maharana et al., 2026).

These mechanisms are of particular interest in the context of neurodegenerative vulnerability. Recent evidence suggests that NPs may contribute to neurodegenerative-like processes, including altered protein aggregation, glial activation, dopaminergic neuronal dysfunction, and behavioural alterations(Araújo et al., 2025, West et al., 2026). However, little is known about how NPs influence neuron–glia homeostasis at the cellular level. Since microglia are key regulators of metabolic support, inflammatory signalling and neuronal stress responses(Brunialti et al., 2025), we investigated whether NP exposure affects microglial energetic fitness and microglia–neuron functional communication.

In this study, 48 h of exposure to polystyrene NPs did not induce overt cytotoxicity, as evidenced by the absence of morphological alterations and unchanged total protein content. Nevertheless, NPs markedly affected the functional interaction between BV-2 microglia and SK-ARE-*luc2* neuronal reporter cells. At low and intermediate concentrations, NPs enhanced the BV-2-mediated neuronal NFE2L2 reporter signal, both under basal conditions and following purinergic stimulation with ATP. In contrast, the highest NP concentration strongly reduced this effect. This biphasic response suggests that microglia may initially engage an adaptive or compensatory program in response to NPs exposure, potentially increasing their ability to support neuronal antioxidant and stress-response pathways. However, when NPs concentration becomes excessive, this supportive function appears to collapse, leading to impaired microglia-to-neuron communication.

Notably, this functional behaviour was closely associated with changes in microglial energy availability. Supportive microglial activity at low and intermediate NP concentrations was accompanied by increased intracellular ATP levels, whereas the loss of communication observed at 200 μg/mL NPs paralleled a reduction in ATP availability. At the same time, MTT conversion was reduced across all NP concentrations, indicating that microglial reductive metabolic activity was already altered even when ATP levels were transiently increased. This divergent relationship between ATP content and MTT conversion suggests that low-to intermediate-level NP exposure does not simply induce a generalized increase in cellular metabolism but rather promotes a selective remodelling of microglial metabolic activity. At high NP concentration, this adaptive profile is lost and shifts toward energetic impairment.

The association between reduced microglial energetic fitness and impaired microglia–neuron communication is particularly relevant in the context of neurodegenerative susceptibility(Brunialti et al., 2021). Similar functional patterns have been observed in experimental models of GBA impairment, a genetic condition associated with increased susceptibility to Parkinson’s disease, in which altered lysosomal–mitochondrial homeostasis compromises microglial metabolism and reduces intracellular ATP levels, as well as microglia’s ability to support neuronal NFE2L2-dependent responses (Brunialti et al., 2025). In this perspective, NP exposure may act as an environmental stressor capable of converging on similar vulnerability pathways, linking particle accumulation, glial metabolic remodelling, and impaired neuronal resilience.

The observation that these effects occur before overt toxicity is particularly intriguing, suggesting that NPs may not simply kill cells but rather silently reshape their functional state. In microglia, this subtoxic remodelling could weaken protective communication with neurons and reduce cellular resilience to subsequent stressors. Over time, such apparently subtle changes may become biologically meaningful, especially in vulnerable brains exposed to additional genetic or environmental risk factors.

Thus, it suggests that NPs can impair microglial metabolic fitness and alter microglia’s ability to modulate neuronal NFE2L2-dependent responses. This may be particularly relevant in conditions of chronic or repeated exposure, where persistent microglial metabolic remodelling could progressively weaken homeostatic and neuroprotective functions (Miao et al., 2023, Sadeghdoust et al., 2024). Although NPs are often considered relatively inert particles, their ability to cross the blood–brain barrier(Licinio et al., 2026), interact with cells, potentially accumulate within intracellular compartments, and perturb essential glial functions may contribute to increased neuronal susceptibility to oxidative or inflammatory stress (West et al., 2026, Araújo et al., 2025).

Overall, these observations highlight the importance of studying nanoplastics toxicity beyond classical cytotoxicity endpoints. Subtle alterations in neuron–glia communication, energetic balance, and cellular resilience may represent early functional warning signals through which apparently inert environmental particles influence long-term brain vulnerability.

## 5. Limitations and future perspectives

This study has some limitations that also define important directions for future research. First, the experiments were performed in immortalized *in vitro* cell models; therefore, validation in primary microglia, neuronal cultures, and *in vivo* models will be necessary to confirm the biological relevance of these findings and to better capture the complexity of neuron–glia interactions in the brain..

Second, the highest NP concentration used here, 200 μg/mL, represents an in vitro high-concentration condition and should not be interpreted as a direct estimate of environmental exposure. Although human studies have reported plastic particles in blood and tissues(Guo et al., 2024, Marfella et al., 2024), including the placenta and brain (de Sousa et al., 2024, Nihart et al., 2025), generally in the ng/mL to low-μg/mL range, quantitative comparisons remain challenging due to differences in analytical methods, particle size ranges, polymer types, and biological matrices. Thus, the dose range used in this study was intended to model increasing cellular NP burden and to identify potential thresholds of functional impairment rather than to reproduce real-life exposure levels. Nevertheless, with appropriate caution, high-burden *in vitro* exposure may provide a simplified experimental framework to investigate cellular consequences potentially associated with repeated or chronic NP accumulation. Future studies should address whether prolonged or repeated exposure to lower NP concentrations similarly affects microglial metabolic fitness, purinergic responsiveness, and microglia–neuron communication. In addition, mechanistic analyses of mitochondrial function, oxidative stress, lysosomal pathways, and inflammatory signalling will be important to clarify how NPs reshape microglial support of neuronal resilience.

## 6. Conclusion

These preliminary findings indicate that nanoplastics can alter microglial metabolic fitness and modulate microglia–neuron functional communication *in vitro*. Low and intermediate NP concentrations appear to trigger an adaptive response characterized by increased ATP availability and enhanced microglia-mediated support of neuronal NFE2L2-dependent stress-response pathways. Conversely, high NP burden impairs ATP-associated responsiveness and reduces microglial support of neuronal stress-response activation. These data support further investigation of nanoplastic effects on microglial homeostatic and neuroprotective functions, particularly in the context of neurodegenerative vulnerability.

## Acknowledgments

The authors are grateful for financial support from the Department of Health Sciences, Linea 2, intramural funding, Università degli Studi di Milano (to E.B. and L.C.). The authors are also grateful for the financial support from PRIN P2022RSWWF, funded by Next Generation EU and MUR (to L.C. and M.P.). We also thank the CRC Nano/microplastics on Environments, Medicine, Ecology, Systems Interaction Studies – NEMESIS, Università degli Studi di Milano. We gratefully acknowledge Dr. Bevilacqua from TÜV Rheinland for his support.

## Author contributions

Conceptualization: E.B., L.C.; methodology: E.B., L.C. Investigation: E.B., C.M.; formal analysis: E.B., A.V.; Funding acquisition: E.B., L.C., M.P.; supervision: L.C., P.C.; writing of the original draft: E.B., L.C., review and editing: A.V., P.C., L.C., M.P.; All authors read and approved the final manuscript.

## Data Availability Statement

The data that support the findings of this study are available from the author upon reasonable request.

